## Supplementary Materials for "Full genome sequence analysis of African swine fever virus isolates from Cameroon"

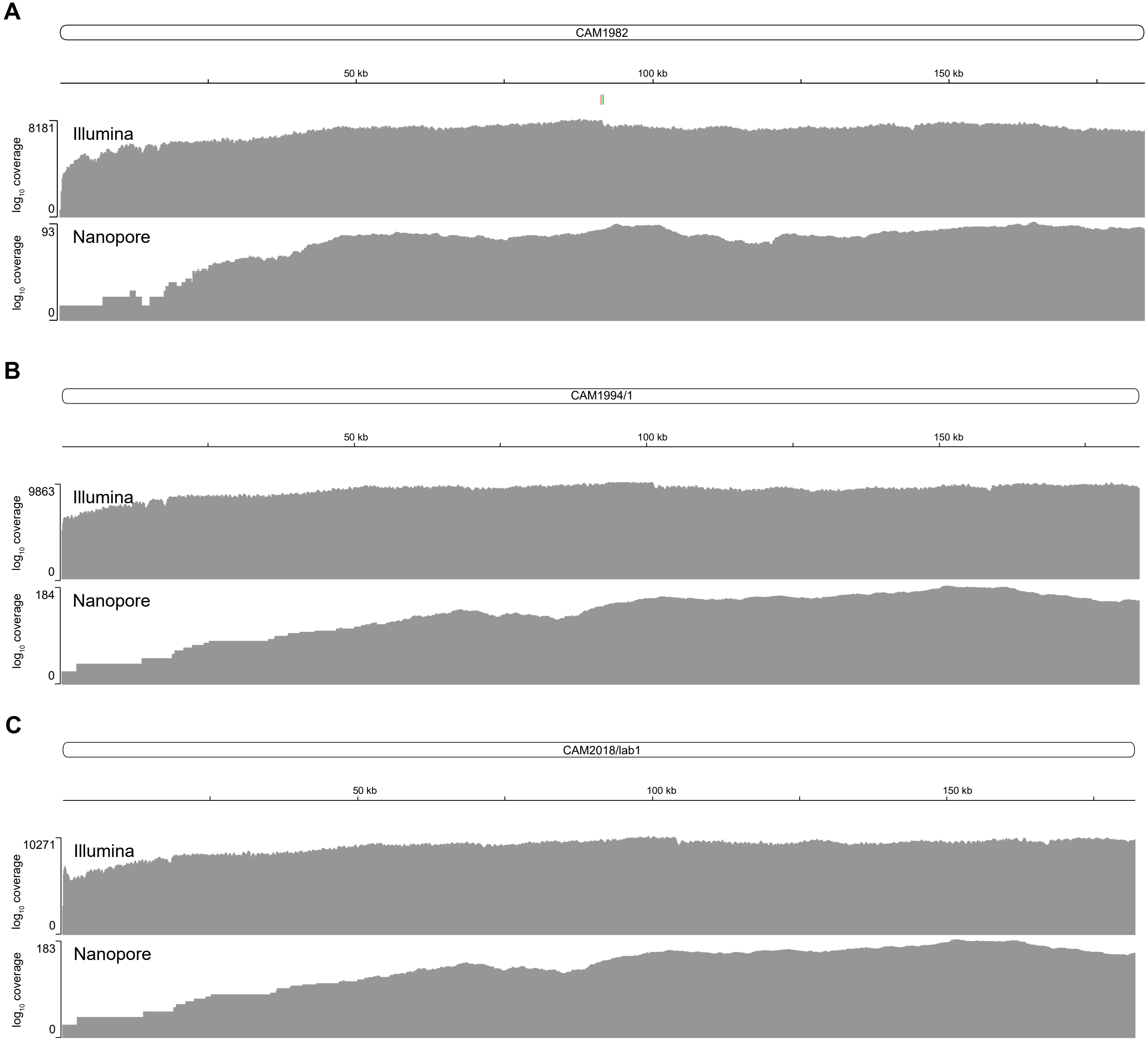


Supplemental Figure S1: Coverage plots. Raw reads from Illumina and Nanopore sequencing runs were mapped back against the assembled genomes of CAM1982 (A), CAM1994/1 (B) and CAM2018/lab1 (C) using Geneious Prime. Plots were displayed using the Integrative Genome Viewer.

| Position in Benin (AM712239) Genome | CAM1982 | CAM1994/1 | CMR2018/lab1 | Gene | Consequence |
|---|---|---|---|---|---|
| 902 | | | G>A | MGF360-1L | P145L |
| 1363 | | | vA | Intergenic homopolymer | |
| 1461 | | | vC | MGF360-2L | Frame shift and truncation. RYKKLCPKIIRWARFII* > RYKNCVRK* |
| 3392 | T>C | | T>C | L83L | K63R |
| 4690 | | | T>C | MGF360-3L | Q59R |
| 6798 | ^CC | ^CC | ^CC | MGF110-11L | Homopolymer. N-terminal change. MKYSWKNGGG > MEKWGGG |
| 7671 | | | vA | MGF110-12L MGF110-13L intergenic, homopolymer | |
| 8187 | | | ^CC | MGF110-13L | Homopolymer. N-terminal change. MGGGDH > MKYSCKHGGGGDH |
| | | | | MGF110-14L | C terminal truncation TWGGVTINNYL* > TWGGG* |
| 9912 | | | G>A | Intergenic | |
| 10113 | vG | ^G | vG | Intergenic homopolymer | |
| 10332 | | ^G | ^G | Intergenic homopolymer | |
| 12086 | | ^G | ^G | Intergenic | |
| 12329 | ^G | ^GG | | Intergenic homopolymer | |
| 12339 | | | G>A | Intergenic homopolymer | |
| 12519 | ^GGG | | ^GGG | Intergenic homopolymer | |
| 14142 | ^GG | ^G | ^G | Intergenic homopolymer | |
| 14367 | vGGG | vGGG | vGGGG | Intergenic homopolymer | |
| 14359 | | | C>T | Intergenic | |
| 14495 | ^GG | ^GG | ^G | Intergenic homopolymer | |
| 14695 | ^G | ^GG | | Intergenic homopolymer | |
| 14873 | C>T | C>T | C>T | J64R | Silent |
| 15735 | | | G>C | MGF300-2R | Silent |

| Position in Benin (AM712239) Genome | CAM1982 | CAM1994/1 | CMR2018/lab1 | Gene | Consequence |
|---|---|---|---|---|---|
| 17207 | | | T>C | Intergenic | |
| 18286 | | ^A | | Intergenic homopolymer | |
| 19549 | vATGTTATAA CC | | | Intergenic repeat (two in Benin, CAM94 and CAM2018) | |
| 24053 | T>G | | T>G | MGF360-12L | M260L |
| 26588 | G>A | | | MGF360-14L | H187Y |
| 28139 | | | G>A | MGF505-2R | R263K |
| 29527 | | | T>A | MGF505-3R | L170Q |
| 34755 | A>G | | A>G | MGF505-6R | N497S |
| 38957 | C>T | | C>T | MGF505-10R | H83Y |
| 39980 | | | C>T | MGF505-10R | L424F |
| 42843 | A>G | | | A151R | K7E |
| 43593 | | | ^T | Intergenic homopolymer | |
| 45261 | T>C | | T>C | A238L | Silent |
| 46995 | | | G>A | A859L | Silent |
| 48371 | | A>G | | A179L | I55T |
| 48639 | | | C>T | Intergenic | |
| 49751 | C>T | | C>T | F317L | V143I |
| 50194 | | | ^A | Intergenic homopolymer | |
| 54233 | T>C | | T>C | F1055L | Silent |
| 61227 | T>C | T>C | T>C | EP1242L | K1073E |
| 63280 | A>G | A>G | A>G | EP1242L | Silent |
| 69507 | | | G>A | EP364R | Silent |
| 70381 | | | G>A | M1249L | Silent |
| 73645 | | | C>G | M1249L | D49E |
| 75103 | | | C>T | M448R | H425Y |

| Position in Benin (AM712239) Genome | CAM1982 | CAM1994/1 | CMR2018/lab1 | Gene | Consequence |
|---|---|---|---|---|---|
| 76117 | | | ^TGCACAAGTGCTTGCACAAGTGCTTGCACAAGTGCC | C44L | 12 aa insertion ASTCASTCASTC |
| | | ^TGCACAAGTGCT | | C44L | 4 aa insertion ASTC |
| 78864 | | | G>A | Intergenic | |
| 80990 | | | C>T | C475L | D146N |
| 81787 | | | G>A | C315R | V104I |
| 82421 | | | ^TTTAAACTAAACG | C315R | Duplication of repetitive sequence, no change to AA sequence |
| 83113 | | | A>C | C62L | S58A |
| 84217 | A>G | | A>G | C962R | Silent |
| 88802 | | | C>T | B962L | R160H |
| 91936 | vGCTTTGGACCGGCCG | vGCTTTGGACCGGCCG | vGCTTTGGACCGGCCG | B169L | Deletion of repetitive sequence. Truncation, removes one copy of three PAGPK repeats |
| 95363 | | | G>A | B602L | Silent |
| 96065 | | | | B602L | CVR, see Table X |
| 96346 | | | | | |
| 96958 | | | ^G | Intergenic | |

| | | | | | |
|---|---|---|---|---|---|
| 101764 | vTGCGTATACTGCCATTGCGTATACTGCCATTGCGTATACTGCCACTGCGTATGCTGCCACTGCGTATACTGCCATTGCGTATACTGCCATTGCGTATACTGCCACTGCGTATGCTGCCAC | | | B407L | Deletion of repetitive sequence, deletion of two of three copies of NGSIR repeat and one of two copies of SGSIR repeat |
| 101765 | | | vGCGTATACTGCCATT | B407L | Deletion of repetitive sequence, deletion of one of three copies of NGSIR repeat |
| 107313 | T>C | | T>C | G1340L | T159A |
| 107316 | | | C>A | G1340L | V158L |
| 109174 | | | C>T | G1211R | Y445 |
| 118226 | | | G>A | CP2475L | A381V |
| 118366 | | | G>A | CP2475L | Silent |
| 121061 | A>G | A>G | A>G | CP530R | I326V |
| 130925 | | | C>T | NP868R | Silent |
| 140155 | | | C>T | D345L | G189S |
| 142963 | T>C | T>C | T>C | P1192R | Silent |
| 145605 | G>A | G>A | G>A | P1192R | H1140R |
| 151564 | ^G | | ^G | Intergenic homopolymer | |
| 160666 | T>C | | T>C | E199L | E127G |
| 160734 | T>A* | | T>A | E199L | Q104H *E199L SNP2 |
| 160793 | C>G* | | | E199L | A85P *E199L SNP1 |
| 163962 | | | | E111R | Silent |
| 164095 | | | G>C | Intergenic | |
| 164257 | | | C>T | I267L | S258N |
| 164364 | A>G | | A>G | I267L | Silent |

| Position in Benin (AM712239) Genome | CAM1982 | CAM1994/1 | CMR2018/lab1 | Gene | Consequence |
|---|---|---|---|---|---|
| 165718 | T>C | T>C | T>C | I226R | Y180H |
| 166580 | A>G | | A>G | I243L | Y4H |
| 167573 | T>C | | T>C | I329L | N183S |
| 168505 | | | ^TCTTCACATTCA | I215L | Duplication of repetitive sequence, additional DECE |
| 169698 | | | ᵛA | I196L | Homopolymer. Truncation at C-terminus, LNLANILNTILCIILIKNV |
| 170250 | | | ᵛA | Intergenic homopolymer | |
| 170745 | T>C | T>C | T>C | DP238L | K99R |
| 171476 | | | ^GGG | MGF360-16R | Extra G at position 88 |
| 171997 | | | ^G | MGF360-16R | 48 C-terminal truncation. |
| 172176 | ᵛCC | | ^C | Intergenic homopolymer | |
| 175204 | | | G>A | Intergenic | |
| 175737 | ^A | | ^A | Intergenic homopolymer | |
| 176084 | | | T>A | I8L | E19D |
| 177456 | | | C>T | Intergenic | |
| 177708 | ^T | | ^T | Intergenic homopolymer | |
| 177718 | | | ᵛA | Intergenic homopolymer | |
| 177827 | | ᵛC | | Intergenic homopolymer | |
| 178121 | | | ^C | MGF360-18R | Frame shift, 59aa deletion. Additional 10 aa changed due to new start codon. Coverage 351. <10% 5Cs |
| 178290 | A>G | | A>G | MGF360-18R | K126E |
| 178869 | ᵛCCC | | ᵛCCC | DP71L | Homopolymer, G4Δ |
| 179486 | | | G>A | Intergenic | |
| 179544 | | | C>T | Intergenic | |
| 181868 | ᵛA | ᵛA | ᵛA | DP60R | Reconstitutes DP60R which is absent in AM712239 |
| 181867 | A>C | | A>C | DP60R | K8Q |

**Supplementary Table S1:** Differences between CAM1982, CAM1994/1 and CMR2018/lab1 genomes and the Benin 1997/1 reference. The positions of substitutions (>), insertions (^) and deletions (ᵛ) relative to the Benin 1997/1 sequence (AM712239) are indicated along with the gene within which the substitution is found, as well as any potential functional consequences.
